# A paradigm to study the learning of muscle activity patterns outside of the natural repertoire

**DOI:** 10.1101/2025.02.13.638098

**Authors:** Ali Ghavampour, Marco Emanuele, Shuja R. Sayyid, Jean-Jacques Orban de Xivry, Jonathan A. Michaels, J. Andrew Pruszynski, Jörn Diedrichsen

## Abstract

The acquisition of novel muscle activity patterns is a key aspect of motor skill learning which can be seen at play, for example, when beginner musicians learn new guitar or piano chords. To study this process, here we introduce a new paradigm that requires quick and synchronous flexion and extension of multiple fingers. First, participants practiced all the 242 possible combinations of isometric finger flexion and extension around the metacarpophalangeal joint (i.e., chords). We found that some chords were initially extremely challenging, but participants could eventually achieve them with practice, showing that the initial difficulty did not reflect hard biomechanical constraints imposed by the interaction of tendons and ligaments. In a second experiment we found that chord learning was largely chord-specific and did not generalize to untrained chords. Finally, we explored which factors make some chords more difficult than others. Difficulty was well predicted by the muscle activity pattern required by that chord. Interestingly, difficulty related to how similar chords’ muscle activity patterns are to the muscle activity patterns required by everyday hand use, and to the overall size of the muscle activity. Together, our results suggest that the new paradigm introduced in this work may provide a valuable tool to study the neural processes underlying the acquisition of novel muscle activity patterns in the human motor system.

## INTRODUCTION

The human motor system possesses a great capacity for learning new motor skills. Professional musicians and elite athletes vividly illustrate the range and complexity of movements that can be attained through repeated practice. Motor skill learning is a multifaceted process (1), involving a range of aspects from temporal sequencing of movements, fine control of speed and force, and more cognitive processes such as action selection and planning.

This paper focuses on a distinct aspect of motor skill learning: that is, the acquisition of entirely novel muscle activity patterns. In fact, many tasks require to synchronously activate specific combinations of muscles that are initially difficult to produce, possibly because they are not part of the existing motor repertoire. For example, the correct hand shape and muscle activity pattern required to play a certain guitar chord (e.g., F# chord) is initially challenging to achieve. Beginners often position their fingers one by one, deliberately using a sequential approach. After training, however, experts can produce the same patterns quickly and synchronously. How novel muscle activity patterns are incorporated in the motor repertoire remains an open question, as well as which neuronal structures are involved in this process.

It is common experience that some muscle activity patterns (e.g., grasping) are intuitive and easy, while others are difficult. One possible explanation is that muscle activity patterns that are needed for everyday hand movements span only a small subspace all possible muscle combinations (2). This subspace may be represented by the nervous system as “muscle synergies”. According to this idea, muscle activity patterns that are far away from these “natural” synergies would be difficult to produce. While there is considerable debate on the neural origin and function of muscle synergies (3–6), the idea that the human motor system has a strong tendency to activate muscles in regular set of patterns is well supported (2). Here we use the term muscle synergy purely at descriptive level, referring to muscle activity patterns that the motor system can generate quickly and synchronously.

Following this definition, learning motor skills such as new guitar chords would require the acquisition of a novel muscle synergy. In the attempt to capture this learning process, we developed a well-controlled task which included “non-synergistic” muscle activity patterns. Our current approach draws from and extends previous paradigms that only assessed independent isometric flexion (7) or extension (8) around the *metacarpophalangeal* (MCP) joint. Here, we combined independent flexion and extension presses, resulting in 242 unique configurations (or “chords”).

In Experiment 1, we verified that despite some chords were initially extremely hard to achieve, all could be successfully mastered with practice. In Experiment 2, participants were trained on a limited set of four chords, showing that performance improvements reflect the specific acquisition of trained muscle activity patterns, rather than general learning. Finally, in Experiment 3, we compared EMG recordings during chord production with those recorded during natural hand actions to show that the muscle activity patterns required for many chords lay well outside the distribution of patterns seen in natural hand use, and that the difficulty of each chord related to its distance from this natural distribution.

## METHODS

### Participants

Fourteen healthy right-handed participants (mean age = 25.1 years, SD = 2.5; seven females) were recruited for Experiment 1. For Experiment 2, we recruited a different group of fourteen healthy participants (mean age = 22.4 years, SD = 3.1; five females), none of whom had previous experience with the task. Finally, ten participants among those who participated in Experiment 1 were also recruited for Experiment 3 (mean age = 25.9 years, SD = 2.1; five females). All participants provided written informed consent before undergoing the experimental procedures. All procedures were approved by the Western University Research Ethics Board (ref 108479) and designed in accordance with the Declaration of Helsinki.

### Apparatus

Participants performed isometric finger presses in flexion/extension directions while keeping their right hand inside a custom-built device (9). The finger box was composed of two five-finger keyboards stacked on top of the other. Foam padding on each key ensured that fingers were comfortably restrained between the upper and lower keys. In addition, the finger box had an adjustable frame by which the space between the upper and lower keys could be adapted to accommodate different hand sizes. Force transducers (Honeywell, FS series, sampling rate = 500 Hz) above and below each key recorded the force exerted by each finger in extension/flexion direction (10 force transducers overall). Finger forces were mapped onto the vertical position of five cursors (white lines, Figure 1b) projected on a screen which moved up and down when the corresponding finger produced force in extension or flexion direction, respectively. The screen projection of the force produced by the ring finger and the pinky was amplified by a factor of 1.5. This adjustment helped maintaining the range of force required to produce chords feasible for all participants; participants were unaware of this adjustment. A grey rectangle marked the resting area (1.2N flexion to 1.2N extension; 0.8N for ring and pinky). Additional horizontal lines above and below the resting area indicated the flexion/extension response areas (2-5N in both directions; 1.3- 3.3N for ring and pinky; see Figure 1).

**Figure 1:**
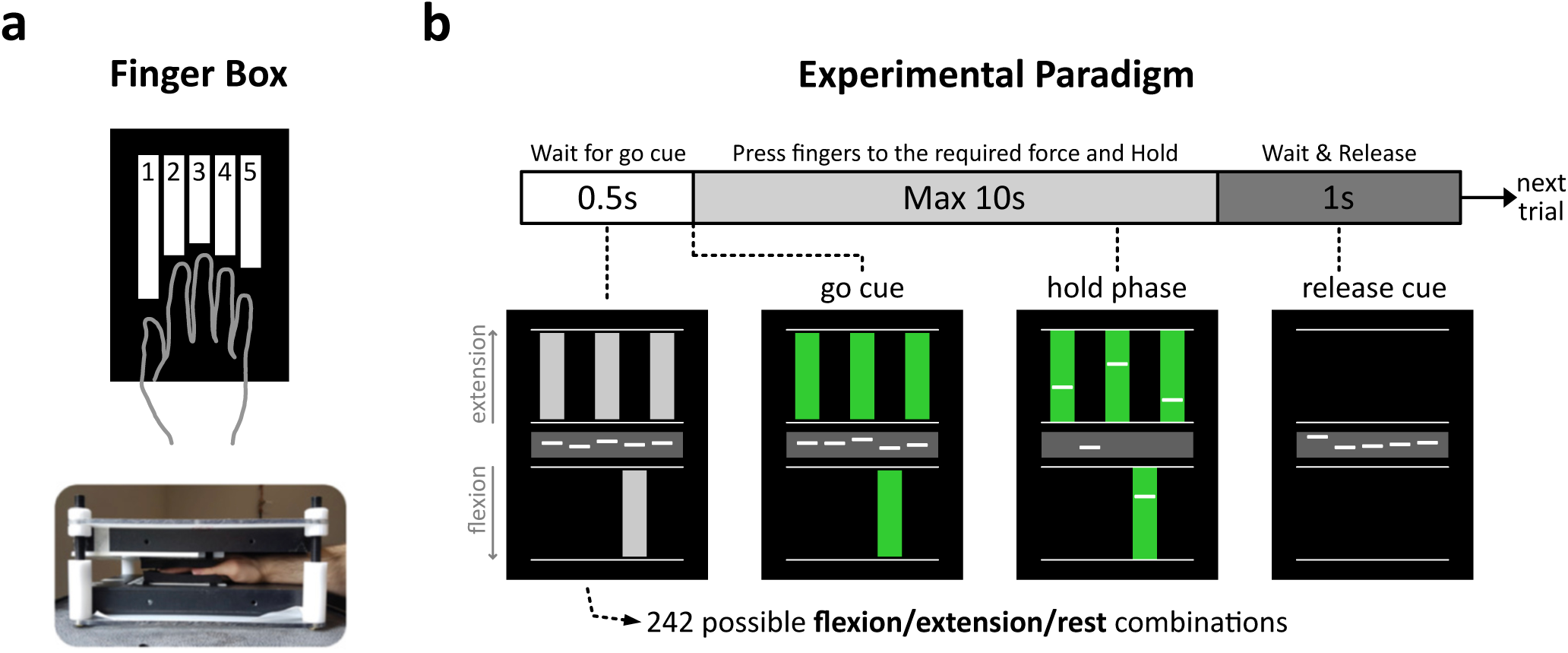
Synergy learning task. **a)** The Finger box measured the isometric flexion and extension forces produced by the 5 fingers. **b)** The task. A spatial visual cue instructed the required forces for chords (here extension of D1, D3, D5, flexion of D4 and holding D2 still). The force level on each finger was indicated by white lines. After the cue turned green, participants had maximally 10s to produce the required force pattern and hold it for 600ms.

### Task

At the beginning of each trial, participants had to keep the cursors inside the resting zone by maintaining their fingers relaxed. The trial began with a short 0.5 second announcement phase, where a chord was presented on the screen with a spatial visual cue (Figure 1b: wait for go cue phase). A total of 242 of chords can be produced by combining finger flexion, extension, or holding still, 242 = 3^5^ − 1 (all fingers still condition). Gray rectangles were presented inside the upper and lower response areas to prompt isometric flexion or extension of the corresponding finger. No rectangle indicated that the finger had to remain still. After 500ms, the rectangles turned green, signalling the go cue. Participants were instructed to produce the required finger forces as quickly and synchronously as possible. Successful chord production was achieved when the chord was held for 600ms. If participants were unable to achieve the required chord within 10s, the trial was marked as unsuccessful. Next, the rectangles disappeared, and participants were instructed to relax their fingers and return to the resting zone. After each trial, a number projected on the screen informed participants whether they successfully achieved the chord (1: successful trial, 0: unsuccessful trial). The following trial started after a 1s inter trial interval. To compensate for slight drifts in force transducers and the posture of the hand, each finger was calibrated by measuring the baseline force while participants were instructed to fully relax their fingers in the finger box.

### Experimental procedures

***Experiment 1*** aimed to characterize the feasibility and difficulty of every possible extension and flexion combination (i.e., chord). Participants practiced all 242 chords over four days, completing 1210 trials per day, organized in 12 blocks (100 trials in the first 11 blocks, 110 trials in the last). Each day included five consecutive trials per chord. The order of chord presentation was fully randomized across days and participants. Participants were verbally instructed to produce quick and synchronous finger presses and encouraged to improve their performance with practice. At the end of each block, participants received feedback on their median execution time (i.e., duration between go-cue and the beginning of the 600- ms hold phase).

***Experiment 2*** aimed to study the learning of specific chords, and the generalization to untrained chords. We selected eight four-finger chords around the median of the mean deviation distribution (see Performance measures below) from days 3 and 4 of Experiment 1. The selected chords were grouped into two sets of four chords. Participants practiced one of the two sets of chords (hereinafter “trained” chords) over five consecutive days, while the other set (“untrained” chords) was only tested on day 1 and day 5. Trained and untrained chord sets were randomly assigned, with seven participants training on set 1 and the other seven on set 2. The task and instructions were identical to Experiment 1. Participants completed 8 blocks of 50 trials each day. Each chord was repeated for five consecutive trials before moving to a different chord and the order of chord presentation was fully randomized across blocks and participants. On day 1 and day 5, trained and untrained chords where randomly intermixed in each block.

***Experiment 3*** aimed to explore the factors that determine the difficulty of chord production. It involved a single session in which we recorded the EMG from hand muscles (see Electromyographic recordings below) both during chord production and natural everyday hand actions. The chord task was structured as in Experiment 1 and 2 and included 68 chords (10 one-finger, 30 three-finger, 28 five-finger chords). Three- and five-finger chords were selected from the lower, middle, and upper thirds of the mean deviation distribution from days 3 and 4 of Experiment 1. This selection procedure ensured that we had chords of different levels of difficulty for each number of fingers. We only selected a subset of the 68 chords to speed up data acquisition.

Participants performed 680 trials (10 trials per chord), divided into 10 blocks, 9 blocks with 70 trials and the last block with 50 trials. After completing the chord task, participants were moved to a nearby experimental setup populated with objects of common use (e.g., calculator, screwdriver, wrenches, door lock and keys, a stylus pen, playing cards, a small whiteboard, an oscilloscope, books, buttons of different shape, a water bottle, zippers). Participants were asked to use their right hand to grasp and manipulate freely these objects for approximately 20 minutes. Importantly, EMG electrode locations (see EMG recordings) remained the same between chord and natural movement recordings. This ensures that natural and chord muscle activity patterns can be described within the same sensor space.

### Performance measures

***Success Rate*** of a chord was defined as the percentage of trials in which the chord was successfully achieved (and held for 600ms) within 10s after the go-cue.

***Execution Time*** was defined as the time interval between the go-cue and the beginning of the 600-ms holding period and reflects how fast chords are achieved.

***Mean Deviation*** quantifies how synchronous fingers are pressed independent of the execution time (7). The force generated by the five fingers in each trial can be described as a five-dimensional trajectory, *F* ∈ ℝ^*T*×5^, where T is the number of time samples between the moment the first finger left the resting zone until the correct force pattern was formed (500Hz sampling frequency). If the fingers are pressed in perfect synchrony, the resulting trajectory will be a straight line (*c* dashed straight line in Figure 2d) from 0 ∈ ℝ^1×5^ (resting force) to the final force pattern, *f*_*r*_ ∈ ℝ^1×5^. To assess finger synchrony, we measured the average Euclidian distance of the produced force trajectory from the straight line:

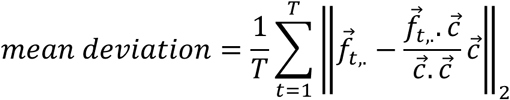

**Figure 2:**
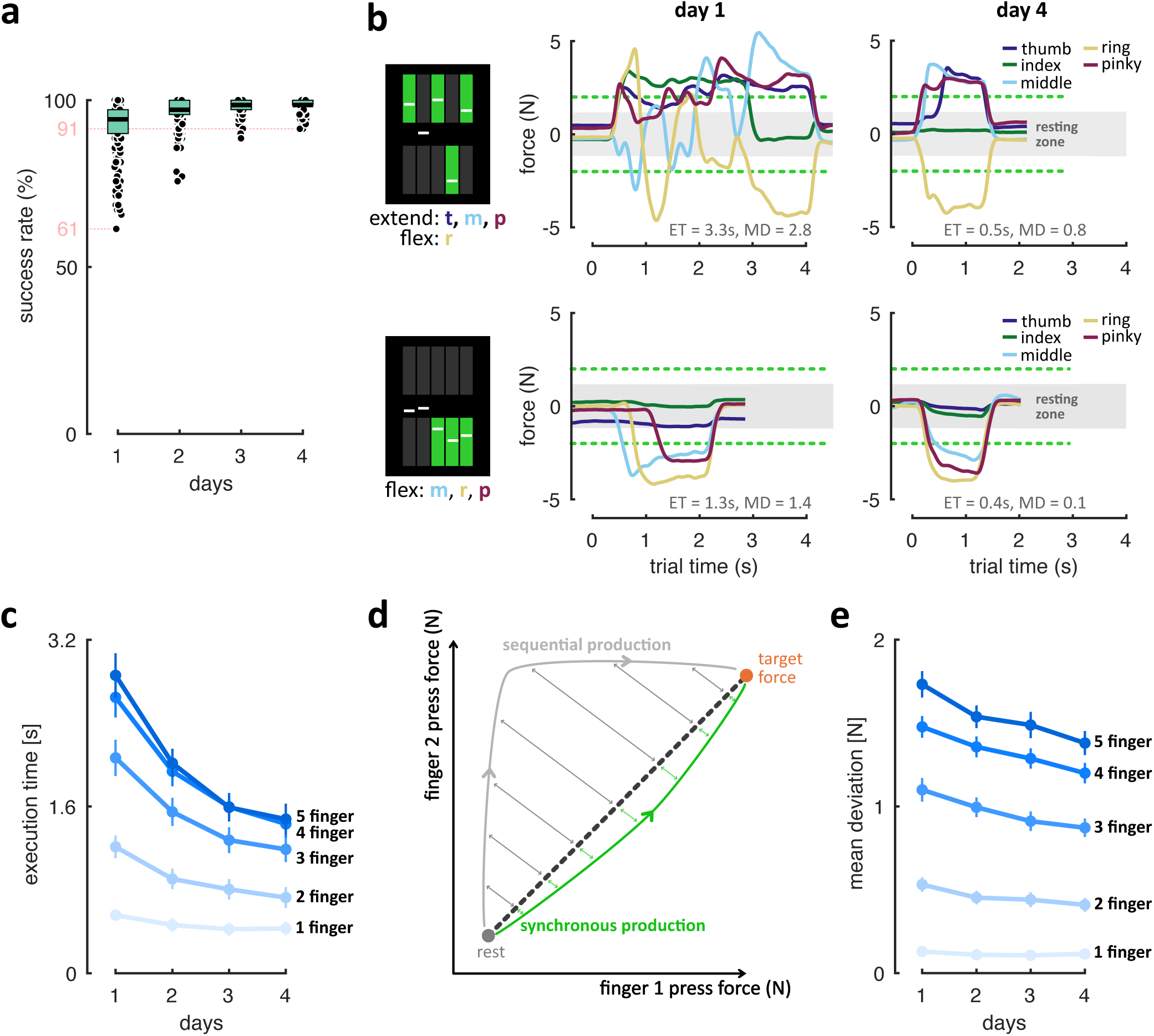
Experiment 1. **a)** Success rate averaged across the 14 participants within each day. Each dot represents a chord. Each box represents the median and interquartile range (IQR). **b)** Example trials of 2 chords from two participants. The green dashed lines denote the minimal required forces in extension (top) and flexion (bottom) directions. Ring and Pinky’s forces are amplified by 1.5x for visualization in this plot. 0s is the go-cue. Execution time (ET) and mean deviation (MD) of the trials are labeled. **c)** Average execution time (s) by day for different number of fingers. Error bars are SEM across participants. **d)** Schematic explanation of mean deviation. Perfect finger synchrony leads to a straight-line trajectory from rest to final force (dashed line). Examples of sequential (gray; high deviation) and synchronous (green; low deviation) force trajectories are plotted. Arrows denote the Euclidean distance from the straight-line. **e)** Average mean deviation by day for chords with different number of fingers. Error bars are SEM across participants.

Where *f*_*t*_, is the force of the 5 fingers at time sample t. Higher synchrony results in a mean deviation closer to 0. Before the calculation, force traces were smoothed using a moving average window with a width of 30 samples (=60ms).

***Response Asynchrony*** was calculated for the chord training study (Experiment 2) where we had enough repetitions of a single chord. To determine the total asynchrony between fingers, we defined the response time of each finger as the time when the derivative of the force crossed 20% of its peak in each trial. In this way, the response time is independent of the baseline force exerted by each finger. The overall response asynchrony between the fingers was then defined as the difference between the response time of the fastest and slowest finger in each trial.

### Electromyographic (EMG) recordings

#### Electrode placement

In Experiment 3, we used a 10-channel surface EMG montage (Delsys, Trigno Research+ System, Trigno Duo Sensors) targeting the muscles acting on the *metacarpophalangeal* joints: the *extensor digitorum communis* (EDC), *extensor digiti minimi* (EDM) and *extensor indicis* (EI), extensors of the thumb (*extensor pollicis brevis* and *longus*), the *flexor digitorum superficialis* (FDS), *abductor pollicis brevis* (APB) and *abductor digiti minimi* (ADM) (Figure 5a: top panel). Raw EMG signals were recorded at 2148 Hz. Electrode locations were selected based on the anatomy of the target muscles and was optimized for each participant by asking them to perform isometric flexion/extension with individual fingers. The electrode was then placed on the surface location where muscle activity appeared maximal through palpation and EMG signal inspection.

#### EMG Pre-processing

The signal from each EMG electrode (sampling rate 2148 Hz, bandwidth 20-450 Hz) was demeaned, rectified and low pass filtered (6^th^ order Butterworth filter, cut-off: 40Hz). The same preprocessing was used for EMG recordings for both chords and natural hand actions.

#### Chord muscle activity patterns

We estimated the muscle activity patterns required for each chord by averaging the preprocessed EMG signals from each channel over the 600-ms holding period and across trials. To account for gain differences across electrode sites, each channel was normalized by dividing by the Euclidean norm of activity across all chords.

#### Distribution of natural muscle activity patterns

After preprocessing, the EMG activity from natural hand actions was averaged within non-overlapping 20ms bins and normalized using the same channel-wise gain factor used for the chord EMG data. The binned normalized data were then partitioned into 10 sets. Each set contained bins sampled 200ms apart by selecting every 10th bin. This approach reduced temporal dependence among samples, which is necessary for the kNN estimator (see below).

### Models of chord difficulty

In the attempt to capture which factors influenced chord difficulty, we designed a set of linear models to predict the mean deviation of each chord using a range of characteristics. Because we wanted to test the predictive power of the muscle activity pattern, we restricted the modelling effort to the 68 chords studied in Experiment 3. The dependent variable was the mean deviation from days 3 and 4 in Experiment 1, resulting in a 68-by-1 vector for each of the 14 participants. To cross-validate, each model was trained on the mean deviation data from 13 participants. We then correlated each model’s predictions with the left-out participant, resulting in 14 correlation values. In the text, we report the predictive performance of models as the mean of these correlation values and the Standard Error of Mean (*SEM*) across participants (*r̅* ± SEM). Models were compared using paired t-test.

#### Noise ceiling

Given that mean deviation was evaluated in the presence of measurement noise, no cross-validated model can reach a correlation of 1 with the data. To estimate the noise ceiling, each participant’s mean deviation vector was correlated with the group mean in which that participant was excluded. The average of the resulted 14 correlation values determined the noise ceiling for the models. The noise ceiling predicts the performance of a group model that captures all systematic differences in chord difficulty across the group.

#### Baseline model

The mean deviation differed substantially for chords with different number of fingers (Figure 2e). Increasing the number of fingers, will increase both the cognitive complexity of the stimulus, and the motoric difficulty. Therefore, the number of fingers was included as baseline in all of our models. We built a design matrix with columns corresponding to an indicator variable (0 or 1) for the number of fingers (three columns for 1-, 3-, and 5-finger chords of Experiment 3).

#### Muscle activity pattern model

This model predicted the difficulty based on the required muscle activity patterns. The design matrix for this model corresponded to the average chord muscle activity patterns across participants (i.e., 68-chords by 10-channels). The average success rate of participants in Experiment 3 was 99.57%, achieving almost 10 successful trials per chord and participant.

#### Force pattern model

This model predicted the difficulty based on the amount and direction (flexion/extension) of the force generated by each finger. The design matrix for this model contained the force generated by each finger in extension and flexion direction averaged over the 600-ms holding period across trials and participants (i.e., 68-chords by 10-forces, 5 regressors for flexion, 5 for extension).

#### Visual complexity model

This model predicted mean deviation based on the visual complexity of the spatial cue used to instruct the chords (Figure 1b). Visual complexity was quantified by counting how many times (in one chord) adjacent fingers received different instructions. For example, all five-finger flexion has 0 transitions, while alternating flexion and extension of the five fingers results in 4 transitions. We built a 68-chords by 5 design matrix, with each column corresponding to an indicator variable (0 or 1) for the number of changes.

### Dissociating muscle activity patterns to magnitude and pattern

#### Magnitude of muscle activity

The magnitude of each chord’s muscle activity pattern 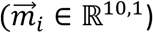 was evaluated by calculating the Euclidean norm 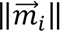, which then was averaged across participants. This resulted in a 68-dimensional vector was used as the group magnitude model to predict the difficulty.

#### Pattern of muscle activity

We evaluated the probability of a chord’s muscle activity pattern, 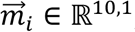, to belong to the distribution of natural muscle activity patterns, using an approach based on *k-nearest neighbour* density estimation. The goal was to estimate the probability of each chord’s pattern to belong to the distribution of natural patterns regardless of the magnitude. Therefore, all muscle activity patterns (both chord and natural distribution) were normalized to unit length (see schematic in Figure 6b). For each normalized chord muscle activity pattern 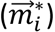, we then ranked the normalized muscle activity patterns from the Natural distribution based on their Euclidean distance to 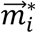. Let *R*_*k,i*_denote the distance from 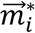 to its k^th^ nearest neighbour in the natural distribution. The kNN density estimator estimates the density for 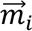 by

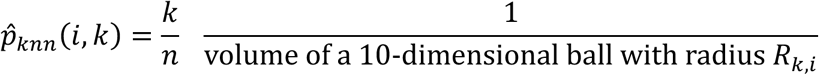

Due to the noisiness of the natural data, instead of choosing a single k, we found more robust estimator for this density by integrating the results across k=0…5. For this we fitted a linear slope passing through 0 and the points defined by 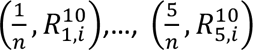 Because the distribution of probability estimates was strongly right skewed, we used the logarithm of this slope as a similarity measure of the chord pattern to the natural distribution. These log-probability estimates (higher values indicated higher similarity) were closer to normal distribution.

To obtain a reliable estimate, the estimation was performed for each chord in each of the 10 natural distributions (see *Forming distribution of natural muscle activity patterns*), and then averaged across them. These estimates were then averaged across the ten participants to derive a group similarity to natural statistics model.

### Alignment of subspaces of the muscle activity patterns for chords and natural actions

The determine the relationship of the space of muscle activity patterns of the chords and the sub-space of natural muscle activity patterns in a cross-validated fashion, we plotted the distribution of natural muscle activity patterns into two non-overlapping splits: the first and third 5-minutes of data were concatenated to create the first split, and the second and fourth 5-minute were concatenated to form the second split. Therefore, each split contained samples of different sets of actions. Principal Component Analysis within each split extracted 10 Principal Components (PC). To cross-validate, the data from one split was projected to the PCs of the other split. The variance explained by PCs then reflects how aligned the probability distribution of natural muscle activities is between different sets of actions. Finally, we projected the 68 chord muscle activity patterns matrix onto the natural PCs to evaluate overall how the chords span the space of the muscle activity. Because natural and chord recordings were performed in the same session without changing electrode locations, the two datasets were directly comparable (see Experiment 3).

## RESULTS

### All chords were biomechanically possible

In Experiment 1, participants were tasked to produce all 242 combination of finger flexion and extension (chords) each day for four days. We first wanted to establish how difficult each chord was, and whether they were biomechanically possible. We therefore counted the number of trials in which participants could produce the chord (success rate). On day 1, the average success rate for some chords was as low as 61% ± 10.8% SEM across participants (Figure 2a, day 1). This indicates that some chords were very challenging, as participants were unable to produce the required force pattern even with 10s allotted per trial. Nonetheless, by day 4, even the most challenging chords were performed with nearly 100% success rate, with the least successful chord reaching a 91% (Figure 2a, Day 4). This indicates that no hard biomechanical limitations prevented chord production and that, with practice, participants could learn the muscle activity pattern required for each chord.

### Both execution time and synchrony improved with practice

The main task goal for participants was to produce the instructed chord as fast as possible after the go- cue. As one performance measure, we therefore used *execution time*, the duration between the go-cue and the moment the correct force pattern was achieved. Execution time (Figure 2c) showed a significant main effect of *day* (repeated measure ANOVA; *F*_3,39_= 58.9, *p* = 1.51e-14) and *number of fingers* (*F*_4,52_= 163.3, *p* < 1e-16), alongside a significant *day* by *number of fingers* interaction (*F*_12,156_= 41.2, *p* < 1e-16). Post-hoc pairwise comparisons indicated significant improvement in execution time from day 1 to 4 for all *number of fingers* (all *t*_13_> 3.06, all *p* < 0.0046). Therefore, chord production became quicker with learning from day 1 to 4.

Our definition of synergy learning demanded that chords should not only be achieved more quickly, but also that they are generated as a single unit with all fingers producing the required force synchronously. Low execution times could conceivably be achieved by producing an asynchronous sequence faster. Therefore, we used the *mean deviation* to measure synchrony between fingers independent of the speed of chord production. A schematic explanation of this measure is shown in Figure 2d. For a sequential trial (gray trajectory; finger 2 is pressed before 1), the force trajectory (in 5- dimensional space) is far away from ideal synchronous trajectory (dashed line). For more synchronous production (green trajectory) the average distance is smaller. Because the distance is averaged across the entire execution time, this measure captures the relative asynchrony of the fingers during execution independent of overall speed.

On day 1, many chords were produced with substantial time gaps between the fingers (see Figure 2b), but performance gradually became more synchronous by day 4. Mean deviation (Figure 2e) showed a significant main effect of *day* (*F*_3,39_= 15.99, *p* = 6.22e-7) and *number of fingers* (*F*_4,52_= 494.9, *p* < 1e-16), as well as a significant *day* by *number of fingers* interaction (*F*_12,156_ = 9.53, *p* = 1.03e-13). Post-hoc pairwise comparisons indicated significant improvement in mean deviation from day 1 to 4 for 2- to 5-finger chords (all *t*_13_> 3.96, *p* < 0.00081), except for 1-finger chords (*t*_13_= 0.91, *p* = 0.19).

### Chord learning is chord-specific

Our results from Experiment 1 indicate that chord production became quicker and more synchronous. Of course, this learning may not indicate the learning of specific, new muscle activity patterns, but rather reflect general improvements (e.g., through familiarization with the task and experimental equipment). In Experiment 2, we therefore quantified how much of the performance improvement was specific to the practiced chords. We selected eight four-finger chords, which had medium difficultly, according to the mean deviation based on Experiment 1. From these, each participant was assigned four *trained* and four *untrained* chords (see methods). The trained chords were practiced for five consecutive days while untrained chords were only tested on days 1 and 5.

On day 1, execution time and mean deviation were not significantly different between trained and untrained chords (paired t-test: both *t*_13_ < 0.57, both *p* > 0.58; Figure 3a). To assess chord-specificity of learning, we measured the difference between the last two blocks of the pre-test to the first two blocks of the post-test (Figure 3b). Execution time and mean deviation of the trained chords improved significantly more for the trained than untrained chords (both *t*_13_> 5.48, both *p* < 1.06e-4; Figure 3b). For untrained chords, the change from pre-test to post-test was not significantly different from zero across participants in either measure (two-tailed one-sample: both *t*_13_< 0.77, *p* > 0.46; Figure 3b: red box). The general improvements were therefore very small compared to chord-specific improvements. This can also be seen in the example chord shown in Figure 3c.

**Figure 3:**
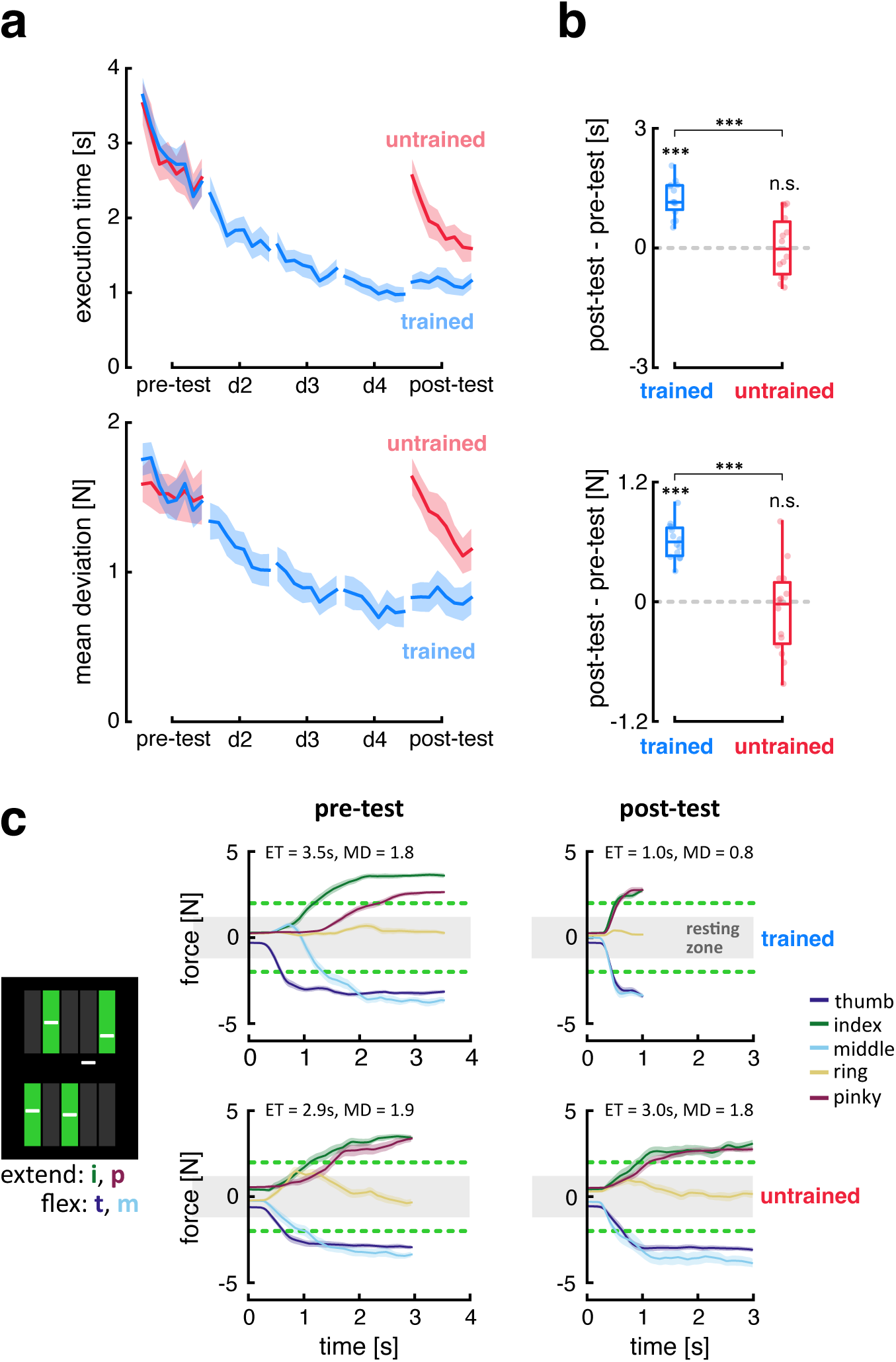
Experiment 2: chord-specific learning. **a)** Average execution time & mean deviation for trained (blue) and untrained (red) chords by practice day. Lines represent the average performance across the 8 blocks with shading representing the SEM across participants. **b)** Improvement of execution time (top panel) and mean deviation (bottom panel) for trained and untrained chords, as measured as the difference between the first two blocks of post-test and the last two blocks of the pre-test. Box plots show the distribution of improvements across 14 participants. Each box represents the median and interquartile range (IQR). Each dot represents a participant. **c)** Finger force traces averaged across all trials of pre- and post-test days. Top panel is when a participant trained on the chord and the bottom panel is for a participant for which the same chord was untrained. The shading represents the SEM across the trials. Force traces are shown from go-cue to the average execution time (*ET̅*) labeled along with average mean deviation (*MD̅*).

### Quantification of remaining asynchrony after practice

While the reduction in mean deviation for the trained chords clearly indicated more synchronous performance, mean deviation did not approach 0 (i.e., perfect synchrony). As mean deviation is a somewhat abstract measure, we also quantified the residual asynchrony between finger presses after learning by looking at absolute time difference between the response of the fastest and slowest finger in each trial (see methods). Mean absolute finger asynchrony on day 5 was 77.64ms ± 20.75 (SEM) for trained and 121.13ms ± 32.37 (SEM) for untrained chords. This range of finger asynchronies suggest that the residual mean deviation may not reflect a deliberate sequencing of the motor commands delivered to the muscles but rather could reflect biomechanical interactions or motor noise.

### Factors determining the difficulty of each chord

A key aspect to make our paradigm suitable to assess synergy learning is to clarify which factors make some chords more difficult to produce than others. This is important to understand what needs to be learned about each chord in order to improve execution performance (i.e., to execute chords faster and more synchronously). The baseline difficulty of each chord is especially relevant for designing training studies based on the current paradigm, for example to match distinct sets of trained and untrained chords. Here we attempted to quantify the role of the cognitive complexity of the stimulus, the patterns of finger forces, and the required muscle activity patterns as determinants of difficulty. As a proxy for chord difficulty, we chose the measure of mean deviation, as it reflects the details of forces exerted by each finger during chord production. Low mean deviation (synchronous performance) suggests that the motor system can easily produce the required muscle activity pattern. To investigate the determining factors of the difficulty of chords, 68 candidate chords were selected based on Experiment 1, including 1-, 3-, and 5-finger chords. The mean deviation measure of these chords, averaged across days 3 and 4 of Experiment 1, spanned a range between 0.045 to 1.870, indicating that they reflected the entire spectrum of very easy to very difficult chords.

***Noise ceiling.*** Before building models to explain differences in difficulty, we established the noise ceiling, i.e., the maximal predictive performance we can expect from a group model (see methods). For this, we determined the average correlation of the mean deviation of the 68 chords from each participant with the group mean (with that subject left out). The average correlation was *r̅* = 0.878 ± 0.0104 Standard Error of Mean (*SEM*) across the fourteen participants of Experiment 1. This high reliability means that the data permits the comparison of different group models of chord difficulty.

***Baseline model.*** In Experiment 1, we found chords involving more fingers had a significantly higher mean deviation. Many factors, including muscle activity and cognitive complexity, increase with the number of fingers. We therefore built a baseline model (see methods) that explains difficulty as an arbitrary function of the number of fingers. This model achieved a predictive accuracy of *r̅* = 0.750 ± 0.0083, indicating that a large component of difficulty can indeed be explained by this general factor. Nonetheless, the baseline model predicted the left-out data significantly worse than the noise-ceiling model (*t*_13_ = 10.89, *p* = 3.33e-8), which indicates that there were easier and harder chords within the groups of three- and five finger chords, and that these differences were consistent across participants. ***Muscle activity pattern.*** This model predicted chord difficulty as a linear function of the muscle activity pattern. To test this idea, in Experiment 3, we recorded EMG from finger flexor and extensor muscles (Figure 4a: top panel), while participants produced the 68 selected 1-, 3- and 5-finger chords. We averaged the rectified EMG during the holding periods (see methods) to estimate the muscle activity required for each of the 68 chords (Figure 4a: bottom panel). We then predicted the mean deviation of chords for each participant in Experiment 1 using a linear model of these 10 EMG channels (see methods). The model predicted the left-out data substantially better than the baseline model (*r̅* = 0.8549 ± 0.0102, Figure 4b, *t*_13_ = 9.37, *p* = 1.89e-7). While the performance was very close to the noise ceiling, it was still significantly lower (*t*_13_ = 10.31, *p* = 1.26e-7).

**Figure 4:**
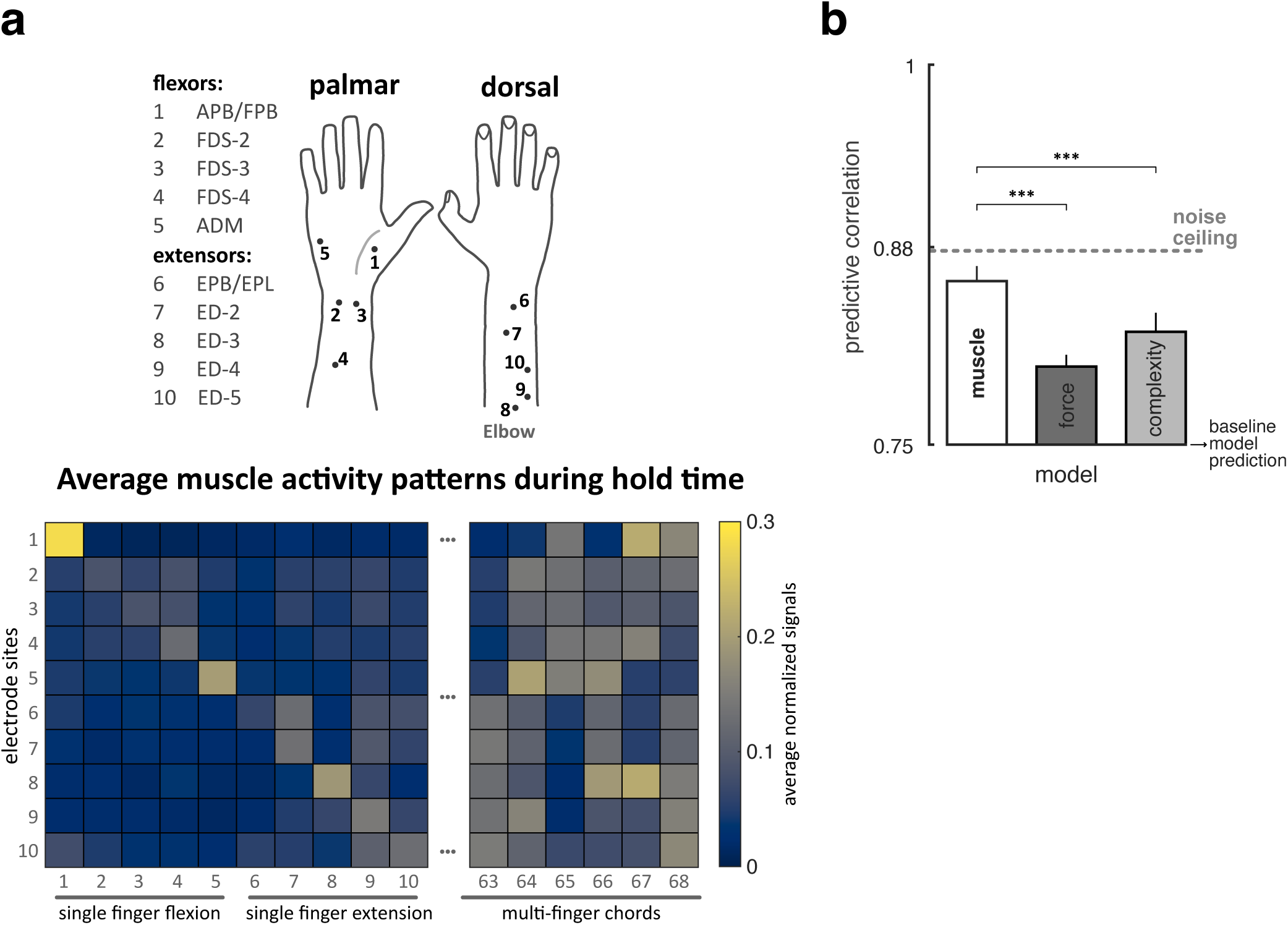
Experiment 3. a) Top panel: Dots indicate the approximate electrode locations on a participant’s dorsal and palmar surfaces of the right hand. The intended muscles are labeled with numbers corresponding to these locations. **Bottom panel:** Average EMG over the 600-ms hold period, averaged across participants. **b)** Cross-validated prediction performance (predictive correlation) of 3 models. The lower limit is the prediction of the baseline model (number of fingers), and the upper limit (gray dashed line) is the noise ceiling. Error bars are SEM across participants.

**Figure 5:**
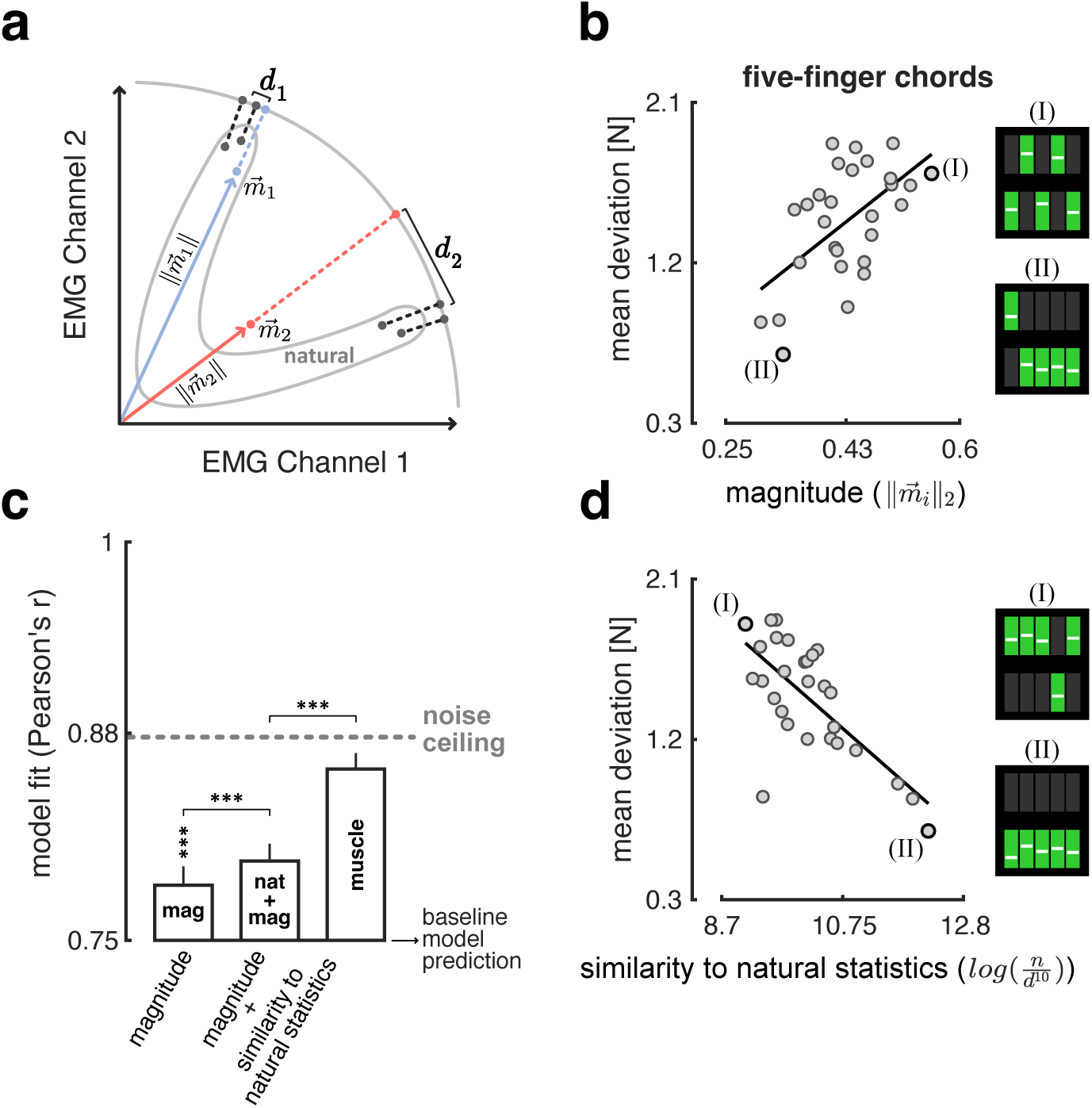
Magnitude and pattern explain the difficulty. **a)** Schematic of evaluating the magnitude (Euclidean norm) and pattern (similarity to natural statistics models) factors. *m*_*i*_s represents two chord muscle activity patterns. The distribution of natural muscle activity patterns is plotted. **b)** Average mean deviation vs. average magnitude for five-finger chords. The chords at the two extremes of both estimates are labeled. **c)** Prediction correlation of magnitude, natural+magnitude, and muscle activity pattern models. The lower limit is the prediction of the baseline model (number of fingers), and the upper limit (gray dashed line) is the noise ceiling. Error bars are SEM across participants. **d)** Average mean deviation vs. average similarity to natural statistics for five-finger chords.

***Force pattern.*** Even though the muscle activity pattern model predicted mean deviation well, the performance needs to be compared against other competing models. We considered that the difficulty might relate to the force direction produced by fingers. For example, chords involving the extension of the ring finger could be more difficult. We therefore predicted the difficulty from the average force pattern required by each chord (1 regressor per finger and direction; see methods). This model outperformed the baseline (*t*_13_ = 9.95, *p* = 1.90e-7) but did not perform as well as the muscle activity patterns (*t*_13_ = 5.64, *p* = 4.02e-5; Figure 4b), suggesting that muscle activity patterns carry more information about the difficulty than force direction.

***Complexity of visual cue.*** Alternatively, we considered that the difficulty can be explained by the cognitive difficulty of processing the visual instructions provided for chords, independent of the amount of muscle activity and the force directions. While we did not have an independent way of measuring “cognitive complexity”, any model based on visual or cognitive factors would predict that two chords with same visual pattern (mirrored across the vertical or horizontal axis) should have the same complexity. To test this idea, we grouped the chords with mirror symmetric visual cues together, resulting in 69 groups among the 242 chords. From these 69 groups of equivalent visual complexity, 37 showed a significant difference (*p* < 0.05) between chords in mean deviation. This number outstrips significantly the expected number of significant results under the Null-hypothesis (5% of 69 = 3.45). This shows that the similarity of the visual cue alone is not sufficient to explain chord difficulty.

To test the performance of this factor alone to predict the difficulty, we designed a model by counting the number of transitions in the spatial visual cue of the chords (see methods). For example, all fingers flexion has zero complexity and alternating flexion, and extension of the fingers has four. Hence, this model is blind to the anatomical information from chords (i.e., which finger should press to which direction) and only captures the overall complexity of the chords. This model performed significantly better than the baseline model (*t*_13_ = 6.51, *p* = 9.84e-6, Figure 4b). However, it performed worse than the muscle activity model alone (*t*_13_= 5.004, *p* = 1.21e-4; Figure 4b), again showing that perceptual or cognitive factors alone cannot fully account for chord difficulty.

### Dissociating magnitude and pattern of muscle activity pattern as determinant of difficulty

We then asked what characteristics of the chord muscle activity patterns determined the chord. A chord’s muscle activity pattern is a point in the 10-dimensional space of muscle activations (Figure 5a: *m*_1_, *m*_2_). The *magnitude* (or length) of the vector captures the overall activation of muscles required for a chord. In contrast, the direction of the vector is determined by the exact pattern of activity across muscles, independent of the magnitude. Here we explored the role of these two factors to the difficulty.

***Magnitude.*** Given that the overall force requirements were identical for all chords involving the same number of fingers, differences in the magnitude of muscle activity may indicate the need for co- contraction of muscles to compensate for the biomechanical interactions between fingers. We therefore predicted that chords that required a higher magnitude should be more difficult. Indeed the magnitude (Euclidean norm) of the chord muscle activity pattern correlated positively with the difficulty for 3-finger (*r̅* = 0.418 ± 0.035; *t*13 = 11.901, *p* = 1.15e-8), and 5-finger chords (*r̅* = 0.383 ± 0.056; *t*13 = 6.86, *p* = 5.73e-6; Figure 5b), and positively but not significantly for 1-finger chords (*r̅* = 0.092 ± 0.077; *t*13 = 1.19, *p* = 0.119). The magnitude model predicted the left-out data significantly better than the baseline model (*r̅* = 0.787 ± 0.012; *t*13 = 4.32, *p* = 4.17e-4; Figure 5c).

***Pattern.*** We also predicted that difficulty should depend on the exact pattern of muscle activities. The motor system is likely to represent some easily producible natural muscle activity patterns. We predicted that if the pattern required by a chord is similar to these natural patterns, it is easier to produce. Within Experiment 3, we therefore also collected EMG while the participants performed naturalistic hand activities. From these data, we formed the distribution of natural muscle activity patterns (see methods). To only account for the pattern, the magnitude was normalized between natural and chord (Figure 5a). Then, we estimated how likely a chord pattern would be under this natural distribution (see methods). These estimates correlated negatively with the difficulty of 1- finger (*r̅* = 0.223 ± 0.068; *t*13 = 3.27, *p* = 3.02e-3), 3-finger (*r̅* = 0.312 ± 0.023; *t*13 = 13.78, *p* = 1.96e-9), and 5-finger chords (*r̅* = 0.451 ± 0.034; *t*13 = 13.08, *p* = 3.71e-9; Figure 5d). The natural statistics model added on top of the magnitude model, significantly outperformed the magnitude alone in predicting the left-out data (*r̅* = 0.801 ± 0.011; *t*13 = 6.897, *p* = 5.45e-6; Figure 5c). This confirmed that, independent of the magnitude, the closeness to the natural distribution of muscle activity patterns can predict the difficulty of a chord.

### Chords span the space of possible muscle activity patterns

Finally, we designed our task such that the muscle activity patterns associated with the set of chords would exhaustively span the space of possible hand muscle activity patterns. It is well know that muscle activity pattern associated with natural hand movements mostly lie within a low-dimensional subspace (2). We therefore wanted to ensure that some of our chords required muscle activation patterns that fell outside of that space. To quantify this, we used Principal Component Analysis to extract the ten Principal Components (PCs) of the natural muscle activity patterns. Using cross-validation we then determined what proportion of the variance was capture by these dimensions using muscle activity recorded during other manual actions (see methods). The results (Figure 6) revealed the expected low-dimensional structure of the natural muscle activity patterns, such that PC 6 to 10 only accounted for 8.92 % of the variance.

**Figure 6:**
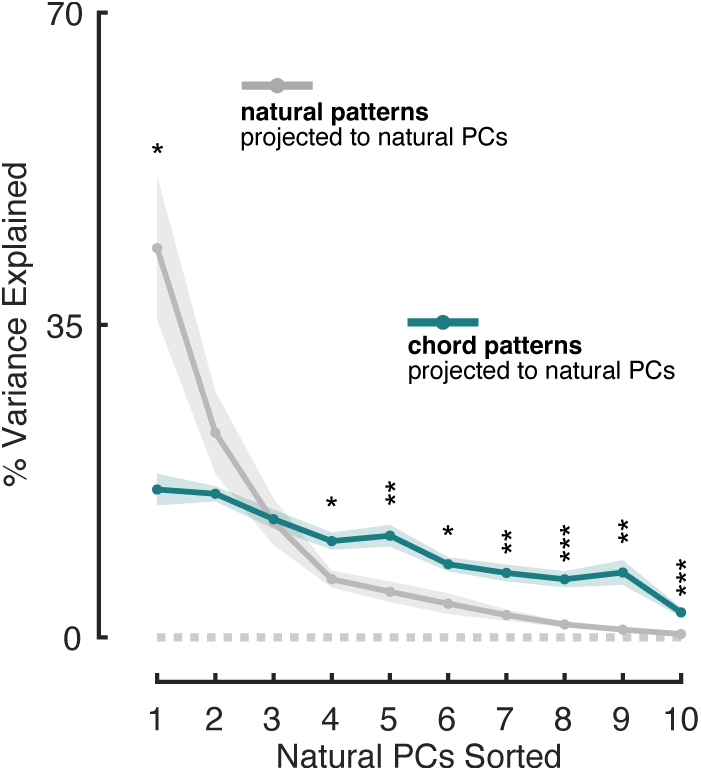
Variance of natural and chord muscle activity patterns projected on every natural PCs. The shading is SEM across participants. PCs are sorted based on the variance they explain in natural patterns. Stars denote the significance of two sample t-test between the variances (*: *p* < 0.05, **: *p* < 0.01, ***: *p* < 0.001).

We then projected the chord patterns onto the natural PCs. This revealed a much flatter distribution of variance, where the least visited five-dimensional subspace by natural accounted for 31.95% of chord variance. This shows that a substantial portion of the muscle activity patterns for the chords required muscular patterns that were rarely produced during natural hand use.

## DISCUSSION

In this paper, we developed a new paradigm to study the acquisition of novel hand muscle activity patterns (“muscle synergies”) that are outside the current behavioral repertoire. Intuitively, this is an important aspect of many skill learning tasks, such as playing the guitar or piano. Our paradigm targets synergy learning by asking participants to perform “chords”— i.e., combinations of isometric finger flexion and extension at the metacarpophalangeal joint. In Experiment 1, we included all the 242 possible chords that can be formed by flexing or extending any combination of 1-5 fingers independently. Some of these hand configurations could be performed immediately and almost effortlessly (e.g., isometric flexion of the index finger) - others were much more challenging. On Day 1, participants failed to produce the most challenging chords in nearly 40% of the trials. This initial low success rate is remarkable, considering that we allowed a 10-second window to achieve chords and repeated each chord 5 times. The allowed time window largely exceeds the usual timescales of everyday voluntary movements and should be sufficient to achieve any force pattern within participants’ motor repertoire. One possibility is that the biomechanical constraints imposed by the intricate interplay of muscles and tendons may prevent the execution of some chords. Tendons and ligaments show quite substantial inter-subject variability (10–12); thus, it was not clear a-priori that all individuals would be able to produce all chords. However, after four days of practice, all 14 participants achieved all chords with nearly 100% success rate. This suggests that all 242 hand configurations are biomechanically possible within the force range required by the paradigm. Therefore, the most challenging chords likely represent genuinely novel hand configurations, which the untrained motor system simply cannot produce. The successful production of these chords after practice therefore suggests the acquisition of new neural representations supporting their execution.

Chords can be achieved by two different strategies: One is to produce the isometric force of each finger sequentially until the desired configuration is reached. This strategy simplifies the problem of controlling different muscles synchronously. Once the required force for one finger is achieved and the corresponding muscle pattern stabilized, another finger is activated, gradually forming the full chord configuration. Indeed, some subjects chose this strategy in the initial part of their training. The improvement in mean deviation across training sessions, however, shows that participant did not improve their execution time solely by producing the same sequence faster. Instead, participants activated the relevant muscles more synchronously. This second “synergistic” strategy suggests that participants formed new chord representations, which allowed them to recruit the corresponding muscle activity pattern both rapidly and synchronously.

This conclusion is also supported by the results of Experiment 2. Here, participants practiced only four four-finger chords. Their performance was assessed before and after training on these four trained chords, as well as on a separate set of untrained chords. Both execution time and mean deviation improved in a chord-specific fashion, with only marginal improvement on untrained chords. This suggests that general learning (e.g., faster processing of the visual stimuli prompting the chords; improved finger individuation) plays a relatively small role in our paradigm. Rather, the results suggest that chord learning mostly occurs through the formation of specific (sensori)motor representations of trained chords.

Yet, it remains unclear where chord-specific learning occurred along the continuum of visual, cognitive and motor processes required for chord execution. Our results do not allow a definite answer. However, some initial insights can be gleaned by assessing which factors make some chords more difficult than others in the first place. Clearly, chords become more challenging as the number of involved fingers increases. However, the number of fingers alone does not fully capture the systematic differences in mean deviation across chords. To determine the most likely explanation for these differences we compared a range of different models: Overall, a model based on the muscle activity patterns outperformed one based on the force pattern and on visual complexity. This suggest that some (if not all) of the chord difficulty is due to the lack of suitable muscle synergies, rather than due the cognitive or perceptual load of interpreting the complex visual cue or monitoring the produced forces via visual feedback.

Mean deviation also scaled with both the magnitude of muscle activity and the similarity of chord muscle activity patterns with those occurring during every-day actions. The latter finding provides evidence that chord performance is indeed constrained by existing synergies: Chords requiring muscle activity patterns similar to those used in everyday motor tasks (e.g., precision/power grip) are easier to achieve compared to those absent in the natural hand repertoire. In other words, learning novel muscle activity patterns may require expanding the current repertoire of motor representations beyond the limited set of existing synergies.

In a related set of studies, participants were asked to acquire an arbitrary mapping between muscle activity or joint kinematics of the upper limb and the 2-dimensional position of a visual cursor (13–17). To some degree, this paradigm(s) requires to learn novel muscle activity patterns. On the other hand, participants were free to choose any pattern in N-dimensional EMG or kinematic space that would map to the same 2-dimensional cursor movement. With only 2 controlled dimensions, it is likely that they chose muscle activity patterns that were closer to their natural repertoire. A key advantage of our paradigm is that it encompasses independent flexion and extension across the five fingers, forcing participants to acquire specific muscle activity patterns, “non synergistic” ones.

The new paradigm introduced in this work is designed to be used in future research addressing the neural correlates of synergy learning with different techniques, including non-invasive brain stimulation and neuroimaging. In this regard, one important question is *where* new synergy representations are formed along the motor hierarchy – from action selection to execution. Previous work suggests that, in sequence learning, new skill representations are not purely motoric but operate at a more abstract level, controlling the sequence of individual movements rather than directly encoding the corresponding muscle activity patterns (18). According to this view, sequence production may rely on the sequential recombination of existing muscle synergies. In support of this idea, a recent study showed that plastic changes accompanying sequence learning mostly occur in high-order motor cortical areas, while being almost absent in the primary sensorimotor cortex (19, 20). Chord learning may share similar principles. Novel muscle activity patterns may be achieved through high-level skill representations combining existing muscle synergies. Alternatively, chords may be achieved through the formation of new motor representations at lower hierarchical level, such as in M1 or spinal cord.

A second important question is *how* multi-finger hand configurations are represented across motor cortical areas. Do multi-finger representations reflect the linear combination of single-finger activity, or do they involve completely orthogonal representations? Current findings are inconclusive. Intracortical recordings performed in the human premotor cortex support the pseudo-linear hypothesis (21). On the other hand, fMRI activity during single- and multi-finger sensory stimulation suggests that the activity patterns for the individual fingers combine predominantly linear in area 3b but show a rich and non-linear behavior in M1 (22). Our task now offers ample opportunity to address these problems in future studies by comparing fMRI activity for single- and multi-finger chords after practice.

## Acknowledgement

We thank Thu Mai for help with data collection on Experiment 2.

## AUTHOR CONTRIBUTIONS

<u>Ali Ghavampour</u>: Conceptualization, Data acquisition, Methodology, Analysis, Writing – original draft, review and editing; <u>Jörn Diedrichsen</u>: Conceptualization, Supervision, Methodology, Writing – review and editing; <u>J. Andrew Pruszynski</u>: Conceptualization, Supervision, Methodology, Writing – review and editing; <u>Marco Emanuele</u>: Data acquisition, Analysis, Writing – review and editing; <u>Jean-Jacques Orban</u> <u>de Xivry</u>: Conceptualization, Methodology, review and editing; <u>Jonathan A. Michaels</u>: Methodology, review and editing; <u>Shuja Sayyid</u>: Data acquisition, Analysis.

